# Cotranslational folding of a pentarepeat β-helix protein

**DOI:** 10.1101/255810

**Authors:** Luigi Notari, Markel Martínez-Carranza, Jose Arcadio Farias-Rico, Pål Stenmark, Gunnar von Heijne

**Affiliations:** Department of Biochemistry and Biophysics Stockholm University, SE-106 91 Stockholm, Sweden; Science for Life Laboratory Stockholm University, Box 1031, SE-171 21 Solna, Sweden

**Keywords:** pentapeptide-repeat protein, beta-helix, cotranslational folding, X-ray structure, *Clostridium botulinum*

## Abstract

It is becoming increasingly clear that many proteins start to fold cotranslationally, before the entire polypeptide chain has been synthesized on the ribosome. One class of proteins that *a priori* would seem particularly prone to cotranslational folding is repeat proteins, *i.e*., proteins that are built from an array of nearly identical sequence repeats. However, while the folding of repeat proteins has been studied extensively *in vitro* with purified proteins, only a handful of studies have addressed the issue of cotranslational folding of repeat proteins. Here, we have determined the structure and studied the cotranslational folding of a β-helix pentarepeat protein from the human pathogen *Clostridium botulinum* – a homolog of the Fluoroquinolone Resistance Protein MfpA – using an assay in which the SecM translational arrest peptide serves as a force sensor to detect folding events. We find that cotranslational folding of a segment corresponding to the first four of the eight β-helix coils in the protein produces enough force to release ribosome stalling, and that folding starts when this unit is ~35 residues away from the P-site, near the distal end of the ribosome exit tunnel. An additional folding transition is seen when the whole PENT moiety emerges from the exit tunnel. The early cotranslational formation of a folded unit may be important to avoid misfolding events *in vivo*, and may reflect the minimal size of a stable β-helix since it is structurally homologous to the smallest known β-helix protein, a four-coil protein that is stable in solution.

## Introduction

With their simple, repetitive architecture composed of a linear array of nearly identical folding units, repeat proteins represent an important paradigm for protein folding studies [1]. Typical repeat proteins are ankyrin repeat proteins, tetratricorepeat proteins, HEAT repeat proteins, leucine-rich repeat proteins, and various kinds of β-helix proteins [2]. In general, the folding of an individual repeat is thermodynamically unfavorable while the interaction between successive repeats is favorable, meaning that a critical number of folded neighboring repeats need to interact in order for the folded state to become more stable than the unfolded state [1]. From an evolutionary point of view it is believed that complex repeat proteins originated by duplication and fusion of smaller identical segments, adding recurrent favorable interactions [3-5].

As for protein folding studies in general, folding of repeat proteins has mainly been analyzed *in vitro* using purified proteins; [6, 7]. *In vivo*, however, proteins can start to fold cotranslationally [8]. For repeat proteins in particular, cotranslational folding, where repeats are added to a growing folding nucleus as they emerge from the ribosome exit tunnel, seems a likely scenario. Still, only a handful of studies have addressed the issue of cotranslational folding of repeat proteins [9, 10].

The pentapeptide repeat protein (PRP) family belongs to the class of β-helix proteins. In PRPs, four pentapeptide repeats form an approximately square repeating unit (a “coil”), and a string of coils form the β-helix [11]. The hydrophobic core inside the β-helix contains conserved Phe and Leu residues, while polar and charged residues decorate its surface, in many cases mimicking a DNA double helix [12]. Here, we analyze the cotranslational folding of a PRP from *Clostridium botulinum* (PENT), a 217-residue polypeptide that forms a highly regular eight-coil β-helix, homologous to the Fluoroquinolone Resistance Protein MfpA from *Mycobacterium tuberculosis* [12, 13]. Using a force-sensing assay based on the SecM translational arrest peptide (AP) [14], we find that an early folding transition takes place when approximately four N-terminal coils have emerged from the ribosome exit tunnel, *i.e*., when only about half the protein has been synthesized; notably, a homologous four-coil PRP from *Nostoc punctiforme* [15] is stable in solution. A second folding transition is seen when the entire PENT domain emerges from the exit tunnel. Since incompletely folded β-helix proteins are prone to aggregation [7], the early cotranslational formation of a folded four-coil unit may be important to avoid misfolding events *in vivo*, and may reflect the minimal size of a stable β-helix.

## Results

### PENT forms a highly regular β-helix

To provide a structural context for our cotranslational folding studies, we first determined the crystal structure of the *Clostridium botulinum* pentapeptide repeat-containing protein PENT (UniProtKB A0A0M0A2X5; see Supplementary Table 1 for refinement statistics and Supplementary Fig. S1 for B-values). PENT crystallizes as a dimer and adopts a right-handed quadrilateral β-helix fold, Fig. 1a, first observed in the *M. tuberculosis* MfpA PRP [13]. Four parallel β-sheets constitute the sides of the square-shaped helix, and its diameter varies between 20 to 30 Å through the squared shape of every coil. The parallel nature of the β-sheets gives a left-handed helicity to the β-helix, each coil being slightly offset from the previous one. The N- and C-terminal caps adopt an α-helical fold that is perpendicular to the axis of the β-helix, and seal off its hydrophobic core. A dimer interface is formed between the C-terminal caps of two PENT monomers, with the α-helices oriented perpendicularly to each other. The two monomers are coaxial, forming a 100 Å long dimer.

**Figure 1.**
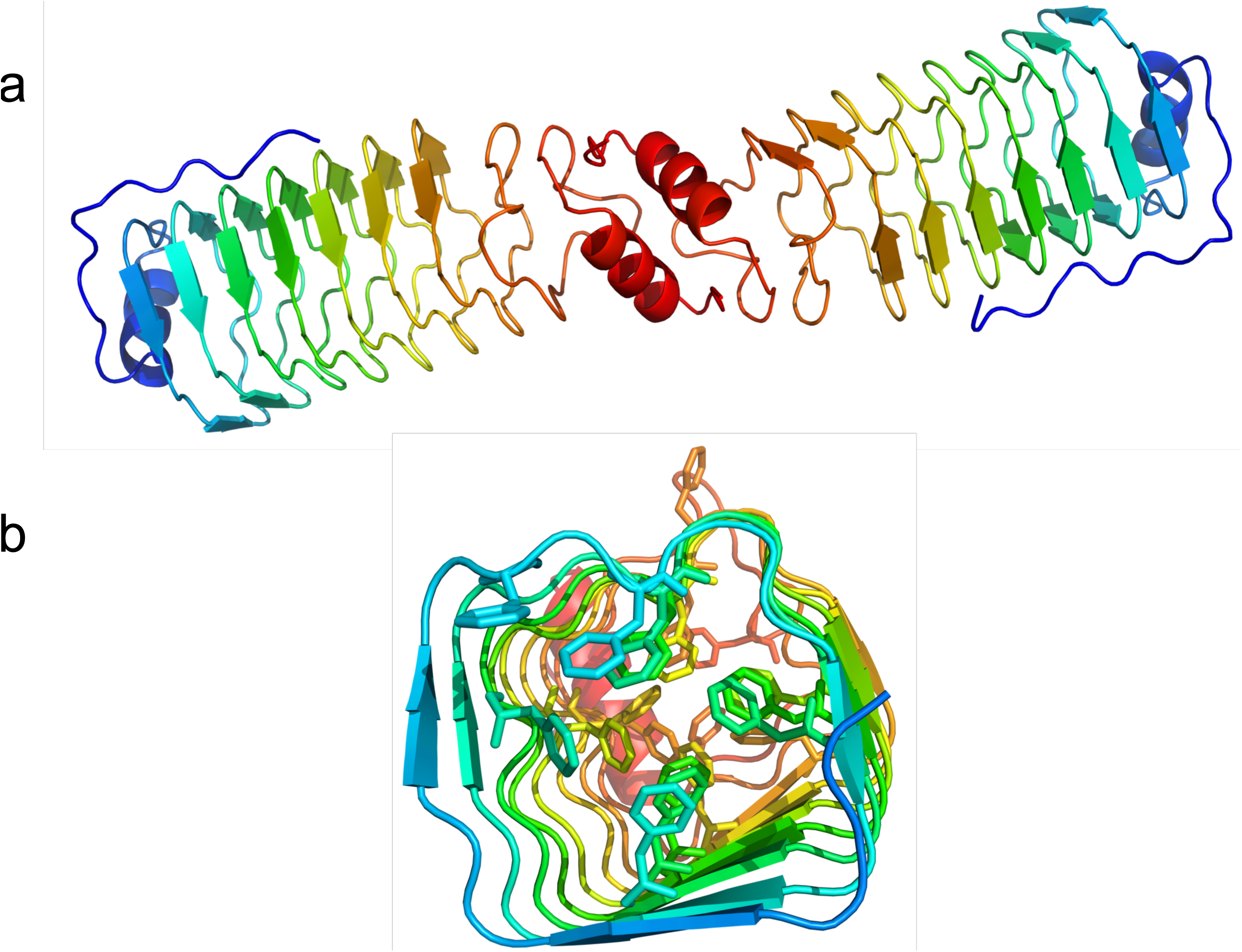
Structural representation of the *C. botulinum* PENT protein. (a) Crystal structure of the PENT dimer. Chains are colored in rainbow from N terminus (blue) to C terminus (red). (b) View along the central axis of the PENT monomer, highlighting the Phe residues in the hydrophobic core. N-terminal residues 1-30 have been removed for clarity.

Each β-helix coil is formed by four pentapeptide repeat units. The central residue of each unit is most often a phenylalanine, and occasionally a methionine, Fig. 1b. These core residues are designated with the *i* position in the repeat, and their side chains point inwards in the β-helix. Residues in the *i^-2^* position also point inwards and constitute the corners of the quadrilateral β-helix. Cysteine and serine residues populate this position most often. The side chains of residues in position *i*^-1^, *i*^+1^ and *i*^+2^ point outwards, and are often populated by amino acids with charged side chains [11, 12]. It is worth noting that no disulfide bonds are formed between any cysteine residues, which might otherwise slow down the folding process. No proline residues are found in the coils, which could otherwise also slow down folding.

### Force-profile analysis of cotranslational folding

Translational arrest peptides (APs) are short stretches of polypeptide that interact with the ribosome exit tunnel in such a way that translation is stalled when the ribosome reaches the last codon in the AP [16]. The stall can be overcome by external forces pulling on the nascent chain [17], and the stalling efficiency of a given AP is reduced in proportion to the magnitude of the external pulling force [18, 19]. APs can therefore be used as force sensors to follow a range of cotranslational processes such as membrane protein biogenesis [18, 20], protein translocation [21], and protein folding [14, 22-27].

A schematic representation of how cotranslational protein folding generates force on the AP is shown in Fig. 2a. For short constructs for which there is not enough room in the ribosome exit tunnel for the protein to fold at the point when the ribosome reaches the end of the AP, or for long constructs where the protein has already folded when the ribosome reaches the end of the AP, little force is generated and translation is efficiently stalled. However, for constructs of intermediate length where there is just enough space in the tunnel for the protein to fold if the tether is stretched out from its equilibrium length, some of the free energy gained upon folding will be stored as elastic energy (increased tension) in the nascent chain, reducing stalling. By measuring the stalling efficiency for a series of constructs of increasing length, a force profile can be generated that shows how the folding force varies with the location of the protein in the exit tunnel [14], and hence when during translation the protein starts to fold. The approach has been validated in a number of ways. It has been shown that the main peak in a force profile correlates with the acquisition of thermolysin resistance of the folding protein in an on-ribosome pulse-proteolysis assay [26], and with the appearance of folded protein in the exit tunnel as visualized by cryo-EM [14, 22, 24]. Further, the amplitude of the peak correlates with the thermodynamic stability of the folded protein [26]. Finally, force profiles can be quantitatively reproduced by molecular dynamics simulations of cotranslational protein folding [14, 24], and are affected by changes in the size and shape of the exit tunnel in ways expected for cotranslational folding [25].

**Figure 2.**
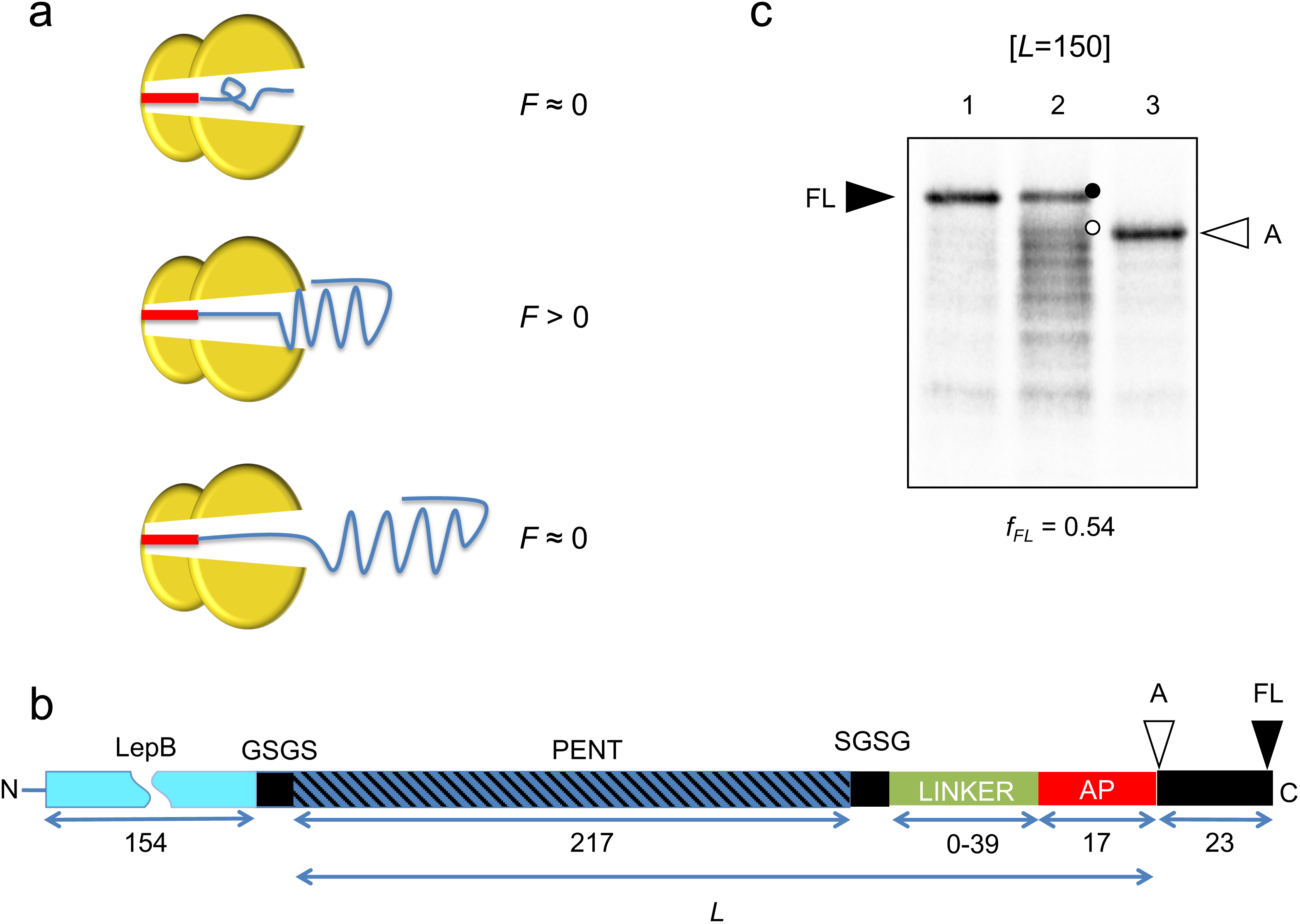
Arrest-peptide based force-measurement assay. (a) Schematic scenario for constructs generating (*F* > 0) or not generating (*F* ≈ 0) pulling force depending on the location of the PENT domain relative to the arrest peptide. (b) The force-generating PENT domain (and C-terminal truncations thereof; blue) is connected, via a variable-length linker (brick red) and an “insulating” SGSG tetrapeptide (grey), to the 17-residue SecM arrest peptide (AP; red). An N-terminal 154-residue segment from the *E.coli* LepB protein (light blue) and a short GSGS segment (grey) is included in all constructs of total length *L* ≤ 216 (where *L* is the number of residues between the N-terminal end of the PENT part and the last residue in the AP) in order to make short PENT constructs conveniently amenable to analysis by SDS-PAGE, and a 23-residue C-terminal tail (also from LepB) is appended at the C terminus in order to make it possible to separate arrested (A) and full-length (FL) chains by SDS-PAGE. The hatched area highlights the β-helix fold within PENT. The lengths of the different parts of the construct are indicated. (c) SDS-PAGE gels showing full-length (FL, black circle) and arrested (A, white circle) species for the [*L*=150] construct (lane 2); bands just below the A band are presumably due to ribosomal stacking. Lane 1 shows a control construct with a Pro⤏ Ala mutation at the C-terminal end of the AP that prevents translational arrest, and serves as a marker for the full-length form of the protein. Lane 3 shows a control construct with a stop codon replacing the C-terminal Pro codon in the AP, and serves as a marker for the arrested form of the protein. The measured *f_FL_* value for the [*L*=150] construct is shown below the gel.

The basic construct used in the force-measurement experiment is shown in Fig. 2b. The C-terminal end of the force-generating moiety (PENT in the present case) is connected via a variable-length linker to the *E. coli* SecM AP [28], which in turn is followed by a 23-residue C-terminal tail (included to allow separation of arrested and full-length forms of the protein by SDS-PAGE). Shorter constructs (up to *L* = 216 residues) also carry a 154-residue segment of the *E.coli* LepB protein at the N terminus, in order to make the protein run in a convenient region of the SDS-PAGE gel. Constructs are translated for 20 min. in the PURExpress^™^ coupled *in vitro* transcription-translation system [29] in the presence of [^35^S]-Met, the radiolabeled protein products are analyzed by SDS-PAGE, and bands are quantitated on a phosphoimager, Fig. 2c. For constructs where little pulling force is exerted on the AP, stalling is efficient and the arrested (A) form of the protein dominates. In contrast, for constructs experiencing a high pulling force there is little stalling and the full-length (FL) form dominates. We use the fraction full-length protein, *f_FL_* = *I_FL_*/(*I_FL_*+*I_A_*) (where *I_i_* is the intensity of band *i* = A, FL), as a measure of the force exerted on the AP [18].

### PENT force profile

To generate the full force profile for PENT, a series of PENT constructs where first the linker and then the PENT-moiety were progressively truncated from the C-terminal end (see Supplementary Table 2 for sequences of all constructs) were translated in the PURE system, and *f_FL_* was plotted as a function of the length *L* of the PENT + linker + AP part, Fig. 3. The force profile has two minor peaks at *L* ≈ 70 and *L* ≈ 115 residues (corresponding, respectively, to PENT truncations at residues ~50 and ~95), and two major, ~20-residue wide peaks at *L* ≈ 145-165 and *L* ≈ 245-265 residues (corresponding, respectively, to PENT truncations at residues ~125-145 and to full-length PENT attached to the ribosomal P-site via a ~30-50-residue C-terminal linker). The latter is at similar tether lengths as seen before for proteins folding in the exit port region of the ribosome tunnel [24, 26], and therefore very likely represents folding of full-length PENT into its native state.

**Figure 3.**
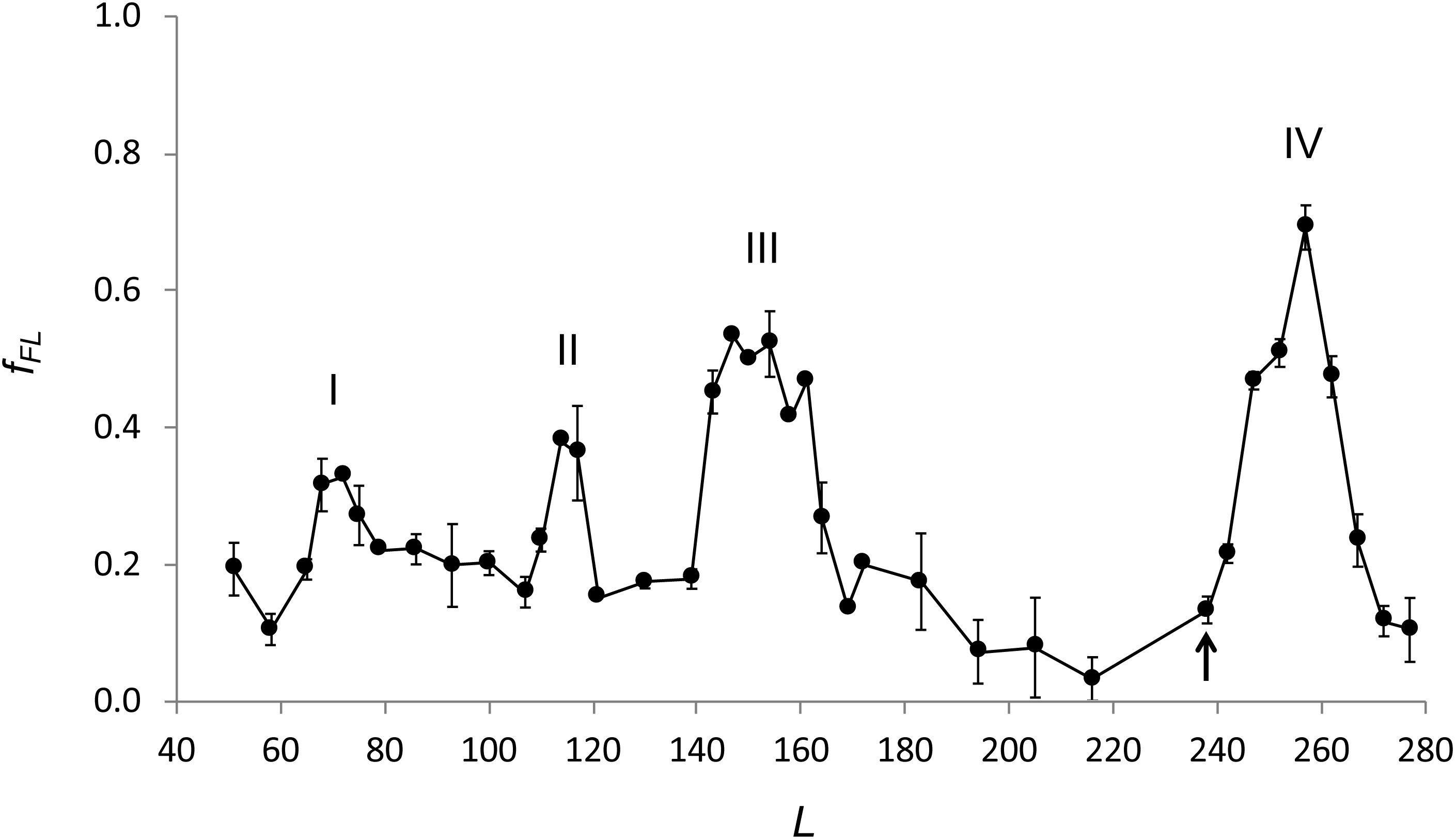
Force profile for the full set of C-terminal truncations of PENT. The four peaks discussed in the text are indicated. The arrow at *L* = 238 residues indicates the shortest construct with the full-length-residue 217 PENT domain (also including the SGSS segment and the 17-residue AP); shorter constructs have C-terminal deletions in the PENT domain and longer constructs have linker segments of 0-39 residues (brick red in Fig. 2b). All experiments were repeated at least 3 times; averages ± SE are shown.

### Effects of mutations in the hydrophobic core of PENT

To ascertain whether the early peaks in the force profile reflect partial folding of the PENT β-helix or may be caused by, *e.g*., minor alterations in the way the nascent chain interacts with the ribosome exit tunnel, we mutated hydrophobic core residues in relevant β-helix coils in the [*L*=114] and [*L*=150] constructs, Fig. 4a. Simultaneous mutation of four core residues in the N-terminal cap and coils 2-3 to Ala (V15A, F18A, F57A, F67A) in the [*L*=114] construct had no significant effect on *f_FL_* (the *f_FL_* value changed from 0.38 to 0.44, data not shown), meaning that peak II most likely does not represent a partially folded intermediate. Peak I is also narrow and of even lower amplitude, and hence is equally unlikely to represent a major folding event.

**Figure 4.**
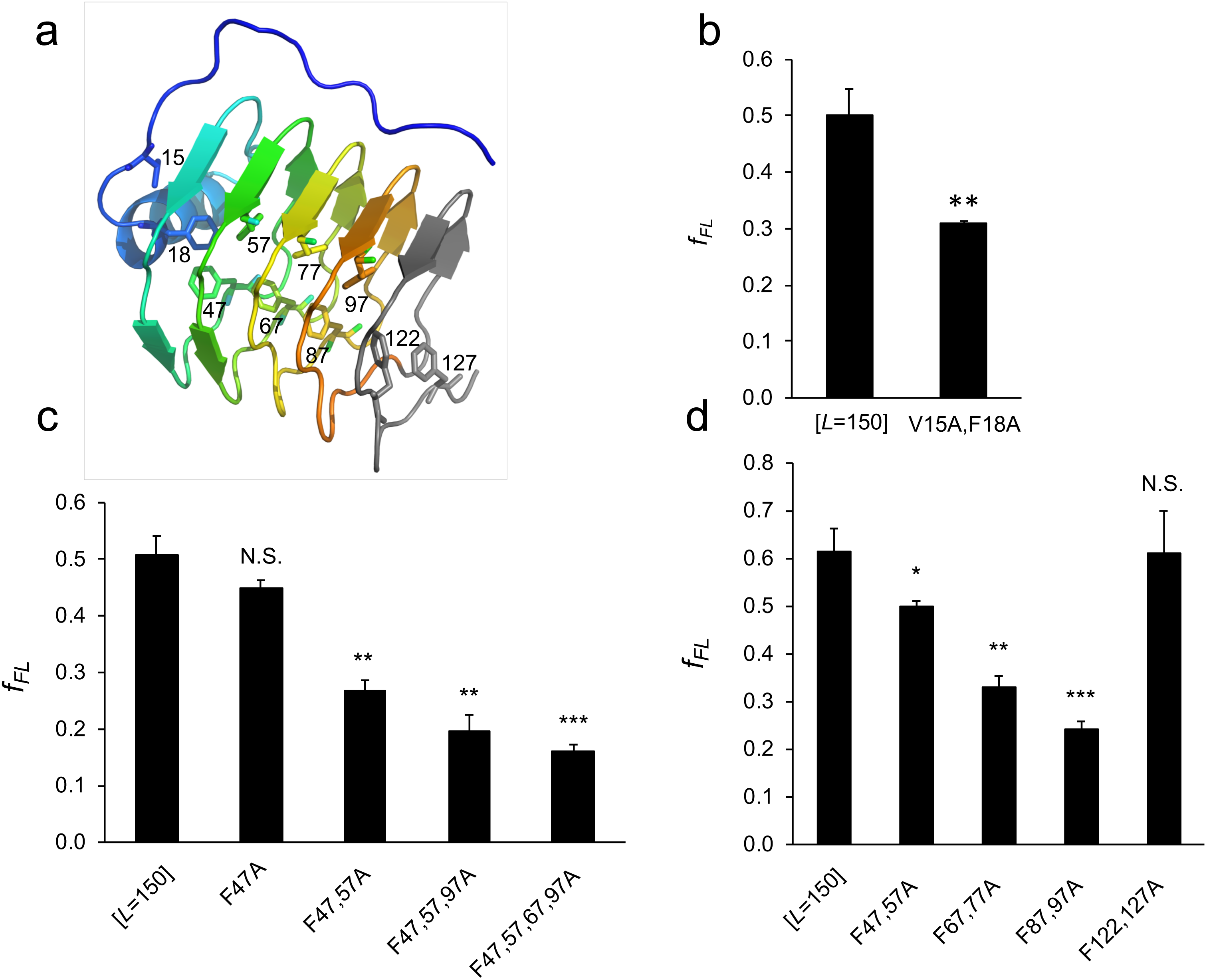
Effects of mutations in the hydrophobic core of the [*L*=150] construct (includes PENT residues 1-129). (a) The N-terminal cap (blue) and 5 N-terminal β-helix coils (indicated by distinct colors) of PENT (residues 1-129), highlighting the hydrophobic core mutations analyzed: Val15, Phe18, 47, 57, 67, 77, 87, 97, 122, and 127. All amino acids were mutated to Ala in a cumulative or paired fashion. (b) Effect of the double mutant V15A+F18A in the N-terminal cap. (c) Effects of cumulative Phe→Ala mutations in the hydrophobic core of PENT. (d) Effects of paired Phe→Ala mutations in the hydrophobic core of PENT. All experiments were repeated at least 3 times; averages ± SE are shown. N.S.: not significant, * *p* ≤ 0.05, ** p ≤ 0.01, *** *p* ≤ Note that the average *f_FL_* value for the [*L*=150] construct (included as a control in all experiments) is somewhat variable, due to different batches of PURE being used for panels b-d.

In contrast, mutations in core residues of the [*L*=150] construct had strong, cumulative effects on *f_FL_*. The double mutation V15A+F18A in the N-terminal cap domain led to a significant decrease in *f_FL_* when compared to the wildtype sequence, Fig. 4b, suggesting that the cap plays an important role in the folding process and stabilizes the folding intermediate [30]. The introduction of multiple F→A mutations within the β-helix coils (F47A, F57A, F67A, F97A) also led to strong reductions in *f_FL_*., Fig. 4c.

To determine the role of each individual β-helix coil, we introduced paired F→A mutations in the hydrophobic core of the [*L*=150] construct, starting in the N-terminal coil and moving progressively towards the C-terminal coil. Paired mutations closer to the C-terminal end of the [*L*=150] construct had stronger effects on *f_FL_*, Fig. 4d, except when placed in the fifth coil (F122A+F127A), where they had no effect. Thus, the fifth coil is not part of the β-helix formed in the [*L*=150] construct, presumably because it remains buried within the exit tunnel.

Finally, to better determine the end of the folded β-helix in the [*L*=150] construct, starting from the C-terminal end of the PENT part (at PENT residue 129) we replaced five amino acids at a time with alternating Gly and Ser residues. Replacement of up to 20 residues had little effect on *f_FL_*, but when the C-terminal 25 residues were replaced there was a strong drop in *f_FL_*, similar to the one seen for the F87A+F97A double mutant, Fig. 5. Thus, the C-terminal end of the folded β-helix in the [*L*=150] construct is located between PENT residues 104 and 109.

**Figure 5.**
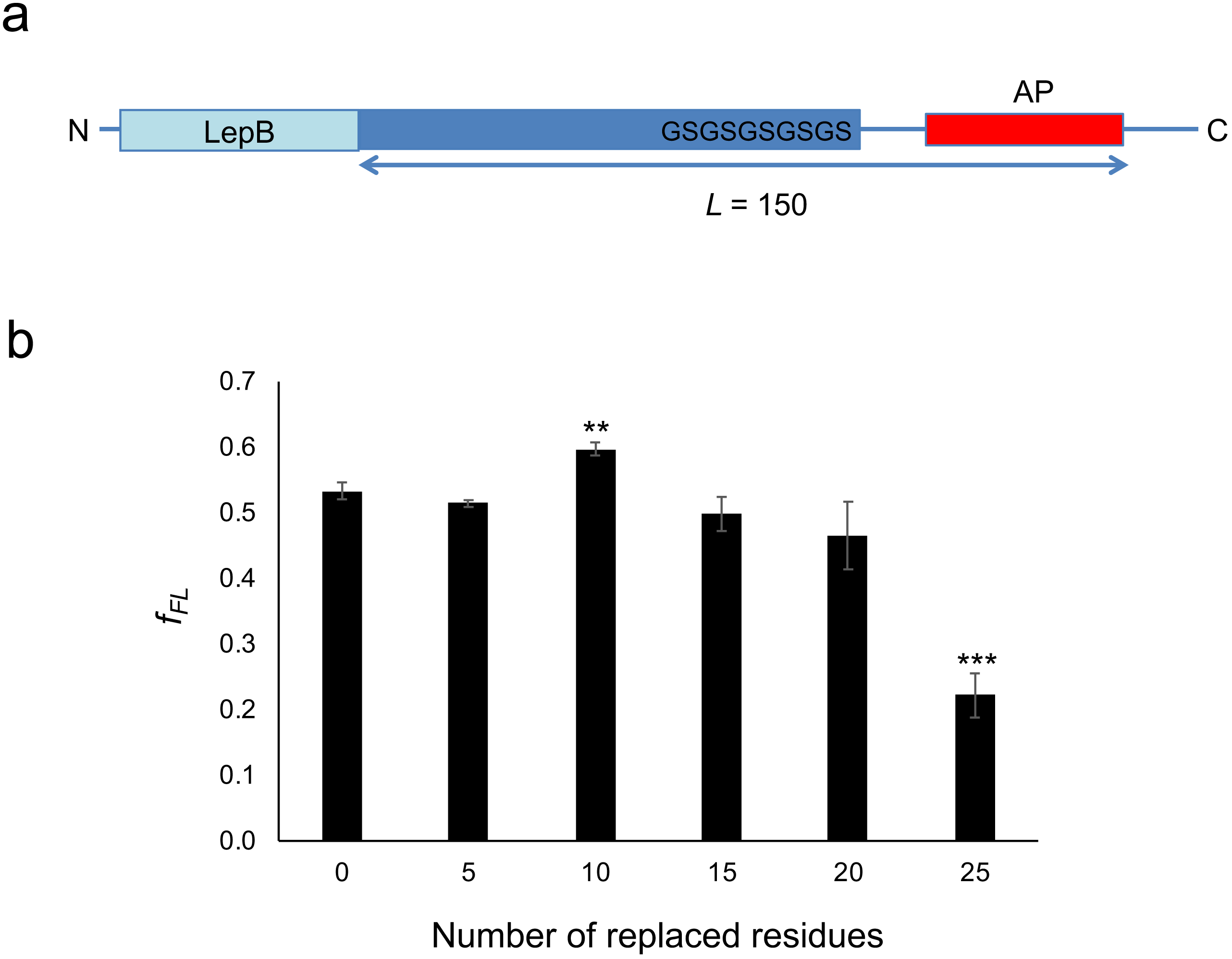
Effect of cumulative replacement of amino acids at the C-terminal end of the PENT moiety in the [*L*=150] construct. (a) Schematic representation of the replacement strategy with alternating Gly and Ser residues, five at a time. (b) Effects of the cumulative replacements. All experiments were repeated at least 3 times; averages ± SE are shown. ** p ≤ 0.01, *** *p* ≤ 0.001.

As a control to ascertain that F→A mutations destabilize full-length PENT, we performed a pulse-proteolysis assay [31] on a [*L*=238/SecM(stop)] construct, which contains full-length PENT and has a stop codon placed at the end of the SecM AP (Supplementary Fig. S2). Construct were expressed in the PURExpress^™^ *in vitro* translation system and subjected to pulse-proteolysis by thermolysin. The [*L*=238/SecM(stop)] construct is fully resistant to thermolysin treatment (average fraction resistant = 0.99). In contrast, [*L*=238 F87A/SecM(stop)] and [*L*=238 F87A+F97A/SecM(stop)] constructs are partially degraded by thermolysin (average fraction resistant = 0.46 and 0.51, respectively). F→A mutations thus both destabilize full-length PENT and perturb the folding of the [*L*=150] construct.

We conclude that the [*L*=150] construct contains a folded β-helix domain that extends from the N-terminal cap to the fourth coil, ending around residue 107±3. The onset of the folding transition is between constructs [*L*=139] and [*L*=143], Fig. 3, meaning that folding commences when PENT residue 107 is 32-36 residues away from the peptidyl-tRNA site (P-site) in the ribosome. This places the C-terminus of the folded four-coil β-helix domain in a similar location in the port of the exit tunnel as we previously determined by cryo-EM for spectrin and titin domains tethered 33-35 residues away from the P-site [22, 24], *i.e*., near a prominent loop in ribosomal protein uL24 at the distal end of the exit tunnel.

### Homology to a stably folded four-coil β-helix protein

We performed a pairwise alignment of profile Hidden Markov Models [32] of the full PENT sequence *vs*. the Protein Data Bank [33] in a search for homologous structures that could explain the folding capabilities of the [*L*=150] construct. The smallest protein that gave a significant hit was the 98 residues long *Nostoc punctiforme* PRP Np275 (PDB 2j8i). Using the Dali server [34], we structurally aligned the PENT structure to 2j8i, Fig. 6, giving scores in support of structural homology (Z-score = 13.3; rsmd = 2.0 Å) [35]. The core Phe residues in PENT mainly align with Leu residues in Np275 (not shown). Strikingly, the structural superimposition between PENT and Np275 covers approximately the same region that was identified as capable of folding by the force-profile analysis (residues 1-97 in Np275 and 10-107 in PENT).

**Figure 6.**
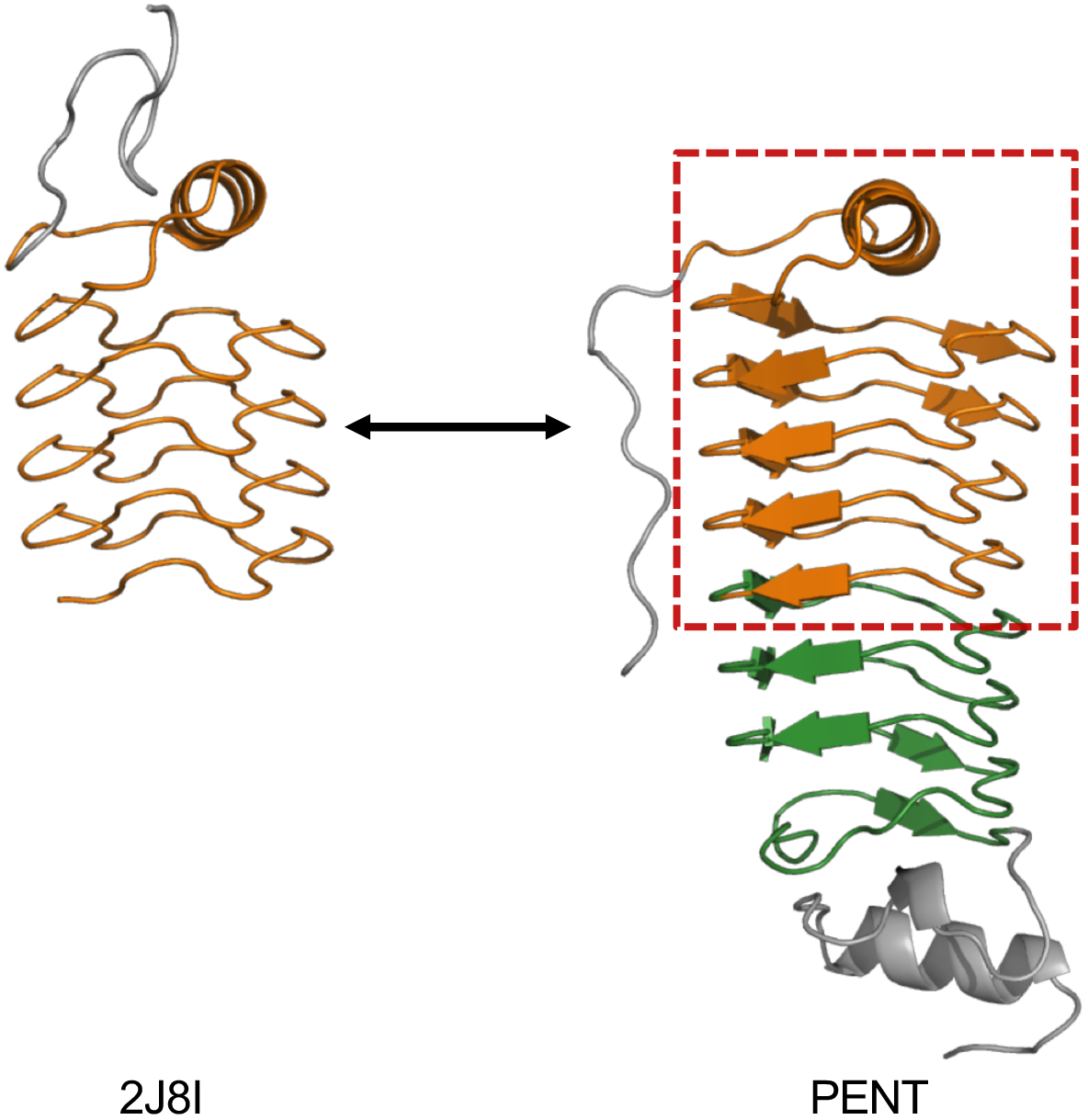
Structural superimposition of PENT and Np275 (PDB 2J8I). The folding unit (PENT residues 1-107) determined by the force-profile analysis is boxed in red.

## Discussion

Despite being obvious candidates to be cotranslationally folding proteins, the only β-helix protein for which there is experimental cotranslational folding data available is the phage P22 tailspike protein, a complex, trimeric protein with a central, 13-coil β-helix domain in the monomer. Using a panel of conformation-specific monoclonal antibodies to analyze the cotranslational folding of P22 tailspike, it was concluded that more than ~3 but less than ~7 coils of the β-helix must be exposed outside the ribosome for the protein to start to fold into a native-like structure [9].

Here, we have determined the structure and analyzed the cotranslational folding of the much simpler and more regular pentapeptide repeat β-helix protein PENT from *Clostridium botulinum.* The structure of PENT is broadly similar to that of MfpA [13], the founding member of the PRP family, except that MfpA lacks the N-terminal α-helix present in PENT. PENT is also structurally similar to the fourth luminal domain of the human synaptic vesicle protein 2C (SV2C), which is a receptor for botulinum neurotoxin serotypes A, D and F from *Clostridium botulinum* [36]. The fact that PENT is present in this bacterium while being similar to the receptor for the neurotoxin it produces is interesting. The fold might hint at the function of the extracellular domain of SV2; it is possible that this domain “mimics” polynucleotides and interacts with DNA- and/or RNA-binding proteins.

Using an arrest-peptide assay that allows us to measure the force generated on the nascent chain by cotranslational folding of PENT, we detect two clear folding transitions: one that involves PENT residues 1 to ~107, *i.e*., approximately the first four coils of the eight-coil β-helix, and one when the full 217- residue PENT domain emerges from the ribosome exit tunnel. It is clear from the force-profile analysis that the first folding transition starts when the C-terminal end of the fourth coil is ~34 residues away from the P-site, at the distal end of the exit tunnel. The subsequent appearance of the remaining coils does not generate appreciable force on the nascent chain until the entire PENT moiety appears in the exit port, suggesting that PENT folds in two steps, where the first step represents folding of the first four coils, and the second represents the addition of the remaining four coils and the C-terminal capping helix to the folded structure.

Interestingly, the four-coil folding unit in PENT (residues 1 to ~107) has significant sequence and structural similarity to the small four-coil PRP Np275 from *Nostoc punctiforme*, suggesting that the four-coil motif may reflect the minimal size of a stable β-helix.

## Materials and Methods

### Enzymes and chemicals

All restriction and DNA modifying enzymes were purchased from Thermo Scientific (Waltham, MA, USA) and New England Biolabs (Ipswich, MA, USA). Oligonucleotides for cloning and mutagenesis were obtained from MWG Eurofins (Ebersberg, Germany). DNA/RNA purification kits were from Thermo Scientific (Waltham, MA, USA). The PUREexpress^™^ cell-free coupled transcription-translation system was from New England Biolabs (Ipswich, MA, USA). [^35^S]-Methionine was purchased from PerkinElmer (Waltham, MA, USA). All other reagents were from Sigma-Aldrich (St. Louis, MO, USA).

### DNA manipulations

All PENT constructs were made from a previously described pET19b-derived plasmid (Novagen, Madison, WI, USA) containing a soluble, non-membrane segment of the *lepB* gene, the small zinc finger domain ADR1a flanked by GSGS, a 39 residues long linker also derived from the *lepB* gene (see Supplementary Table S2 for details), the *E. coli* SecM arrest peptide, FSTPVWISQAQGIRAGP, and a 23-residue C-terminal tail, under the control of a T7 promoter [14]. The GSGS-flanked ADR1a gene was replaced with the 217 residue *M. tuberculosis* PENT gene, using the megaprimer approach. C-terminal truncations of PENT and the *lepB*-derived linker (Supplementary Table S2) were generated by PCR using partially overlapping primers, as previously described [18]. Site-directed mutagenesis was performed to generate constructs with the non-functional FSTPVWISQAQGIRAG<u>A</u> SecM AP as full-length controls for all constructs, to replace the Pro at the end of the AP with a stop codon as arrest controls for all constructs, and on the truncated PENT [*L*=150] and [*L*=161] constructs to introduce Phe→Ala mutations in the PENT hydrophobic core. Cumulative mutations were introduced by site-directed mutagenesis using partially overlapping primers, and mutations of paired Phe residues in the hydrophobic core of PENT were introduced by Gibson assembly [37] using synthetic gene fragments encompassing the mutations (GeneArt Strings, Thermo Scientific, Waltham, MA, USA). All constructs were verified by DNA sequencing.

### Expression in vitro

*In vitro* coupled transcription-translation was performed in PURExpress^™^ (New England Biolabs, Ipswich, MA, USA), according to the manufacturer’s instructions, using PCR products as templates for the generation of truncated nascent chains. Briefly, 1 μl (50 ng) of PCR template and 10 μCi (1 μl) [^35^S]-Methionine were added to a final volume of 10 μl reaction components.

Transcription-translation was carried out for 20 min. at 750 rpm in a bench-top tube shaker at 37°C. Reactions were stopped by adding equal volumes of ice-cold 30% trichloacetic acid, incubated on ice for 30 min., and then centrifuged at 14,000 rpm (20,800 x g) at 4°C for 10 min. in an Eppendorf centrifuge. Pellets were resuspended in 20 μl of SDS Sample Buffer and treated with RNase A (400 μg ml^− 1^) for 15 min. at 37 °C. Proteins were separated by SDS-PAGE, visualized on a Fuji FLA-9000 phosphoimager, and quantified using ImageGauge software V4.23 (FujiFilm Corporation). Analysis of quantified bands was performed using EasyQuant (in-house developed quantification software). Values of *f*_FL_ were calculated as *f*_FL_ = *I*_FL_/(*I*_FL_ + *I*_A_), where *I*_FL_ is the intensity of the band corresponding to the full-length protein, and *I*_A_ is the intensity of the band corresponding to the arrested form of the protein. Experiments were repeated at least three times, and SEMs were calculated.

### Protein expression and purification

PENT was expressed in *E. coli* BL21(DE3) T1R pRARE2, grown in TB supplemented with 8 g/l glycerol and 0.4% glucose, with 50 μg/ml kanamycin and 34 μg/ml chloramphenicol at 37°C. After OD_600_ of 2 was reached the temperature was lowered to 18°C and expression was induced with 0.5 mM of IPTG. Protein expression continued overnight, and cells were harvested by centrifugation (10 min. at 4,500 g) the next morning. Cell pellets were resuspended 100 mM HEPES pH 8.0, 500 mM NaCl, 10% glycerol, 10 mM imidazole, 0.5 mM TCEP. A tablet of Complete EDTA-free protease inhibitor cocktail (Roche) and 5 μl/ml benzonase nuclease (250 U, Sigma) were added to the resuspension, and cells were lysed by pulsed sonication (4 s on/4 s off for 3 min., 80% amplitude in a Vibra-Cell Sonics sonicator). The lysate was centrifuged for 20 min. at 49,000 g and the supernatant was collected and filtered through a 0.45 μm pore size filter.

A 2 ml HisTrap HP column (GE Healthcare) was used for the first purification step. After running the lysate through, the column was washed with IMAC buffer (20 mM HEPES pH 7.5, 500 mM NaCl, 10% glycerol, 0.5 mM TCEP) containing 10 mM imidazole. A second wash was performed with IMAC buffer containing 50 mM imidazole, and the protein was eluted with IMAC buffer containing 500 mM imidazole. The elution fraction was loaded onto a HiLoad 16/60 Superdex 75 column (GE Healthcare) as a second purification step. Fractions were examined on an SDS-PAGE gel, and TCEP was added up to 2 mM concentration before pooling and concentrating with a centrifugal concentrator.

### Crystallization and structure determination

PENT crystals were found in the A6 condition of the PACT premier screen (Molecular Dimensions). In order to optimize these, 150 nl of protein solution was mixed with 50 nl of reservoir solution (0.1 M SPG buffer pH 9.5, 25% PEG 1500) in a sitting-drop crystallization plate. Long thin rod-shaped crystals grew overnight, and were flash frozen in liquid nitrogen after transfer to a drop of cryo-protecting solution (0.1 M SPG buffer pH 9.5, 6.6 mM HEPES pH 7.5, 100 mM NaCl, 10% glycerol, 25% w/v PEG 1500). X-ray diffraction data collection was performed at beamline i03, Diamond Light Source (Oxford, UK). The data were processed using DIALS [38], molecular replacement was performed using Phaser [39] with a model provided by the Phyre^2^ server [40]. The model was built using Phenix Autobuild [41] and Coot [42], and the structure was refined using Refmac5 [43]. Data processing and refinement statistics are presented in Supplementary Table 1. Coordinates for the PENT structure have been deposited in the PDB with the accession code 6FLS.

### Sequence and structure comparisons

The hhpred server [44] was used with default parameter settings (https://toolkit.tuebingen.mpg.de/#/tools/hhpred) to search the PDB database with the full PENT sequence (residues 1-217). The Dali server [34] http://ekhidna2.biocenter.helsinki.fi/dali/ was used with default parameter settings to generate the structural superposition shown in Fig. 6.

### Pulse proteolysis

To assess the stability of full-length PENT protein and mutants thereof, we employed a pulse-proteolysis assay [45]. Full-lenght PENT protein (217 residues) was cloned upstream of a 21 residues long linker composed of alternating Gly and Ser residues followed by the SecM(*Ms*) arrest peptide [46] with the last Pro residue mutated to a stop codon. After *in vitro* synthesis of radio-labelled protein in the PURE system as described above, the reaction was stopped by addition of chloramphenicol (instead of TCA) at a final concentration of 3.3 mM. Each sample was divided in two aliquots of equal volume, one for pulse-proteolysis [31] and the other serving as a non-treated control. Pulse-proteolysis was performed by treatment with thermolysin (buffered in 50mM Tris, 500 μM ZnCl_2_, pH7.0) at a final concentration of 0.75 mg/m for 1 min. at 37°C, under constant shaking at 700 rpm. The same conditions were applied for the non-treated controls with the difference that thermolysin was not included in the buffer solution. The reaction was stopped by addition of 3μl 500mM EDTA (pH 8.4), and protein precipitation was performed by addition of 1:1 volume of ice-cold 10% TCA and incubation on ice for 30 minutes. As described in the previous section, after centrifugation at 14,000 rpm for 5 minutes at 4°C, the supernatant was discarded and the pellet was re-suspended in sample buffer at 37°C for 15 minutes, under constant shaking at 900 rpm. The samples were supplemented with RNase I (670 μg/mL) at 37°C, for 30 minutes, and were subsequently resolved by SDS/PAGE. All pulse-proteolysis assays were performed in triplicates.

## Author contributions

GvH and PS conceived the project. LN, MMC, and JF designed and performed the experiments. LN, MMC, JF, PS, and GvH wrote the manuscript.

## Acknowledgements

We thank the beamline scientists at DLS, Oxford, BESSY, Berlin, ESRF, Grenoble, Max-Lab, Lund and SLS, Villigen for their support in data collection and PSF for protein production, and Dr. Ola Nilsson (Stockholm) for technical assistance. This work was supported by grants from the Knut and Alice Wallenberg Foundation, the Swedish Cancer Foundation, and the Swedish Research Council to GvH, and by the Swedish Cancer Foundation and the Swedish Research Council (2014-5667) to PS.

---

**Supplementary Figure S1.**
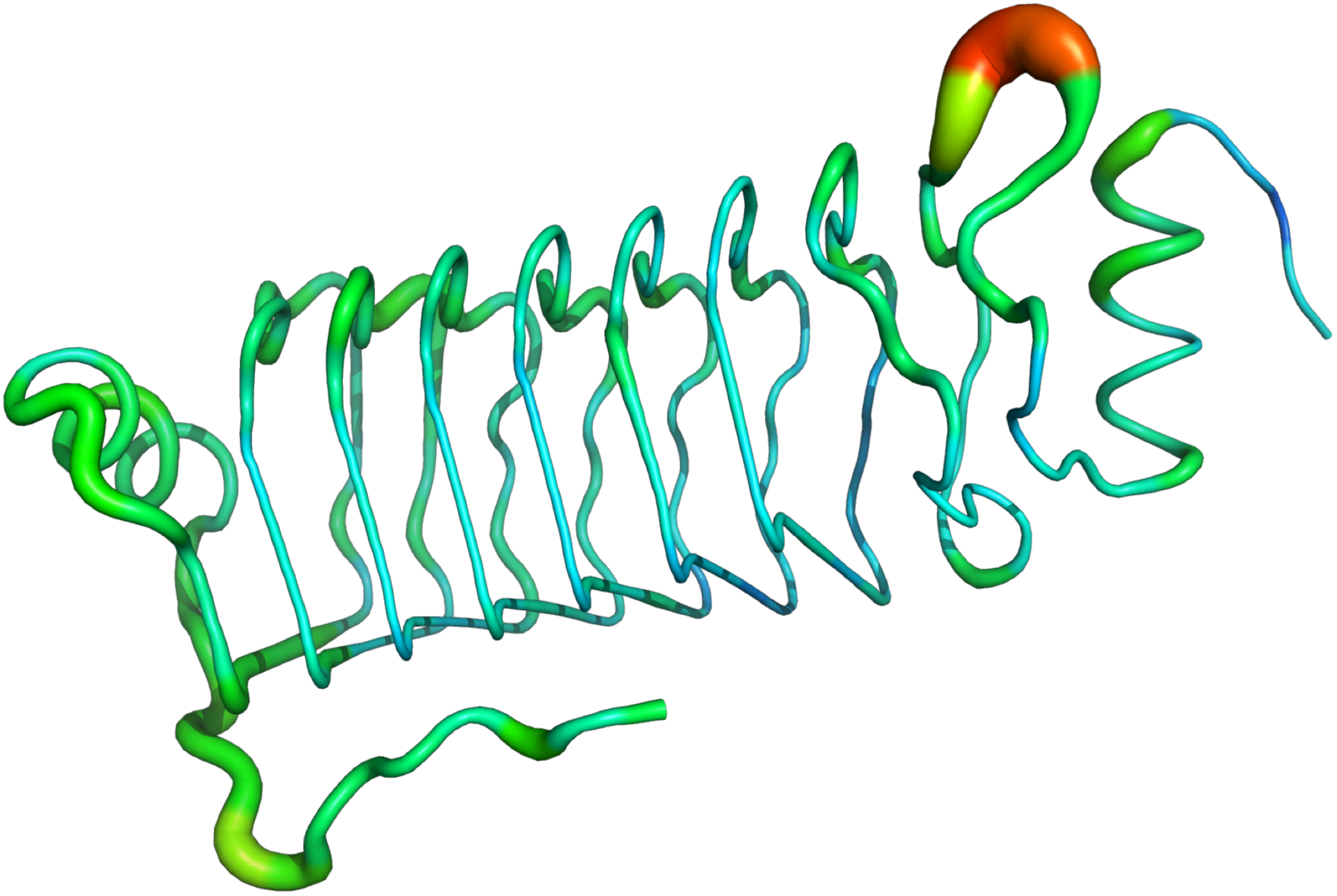
Cartoon putty representation of chain A in the crystal structure of PENT, coloured by B-factors (low B-factors represented in blue and narrow radius, high B-factors in red and wide radius).

**Supplementary Figure S2.**
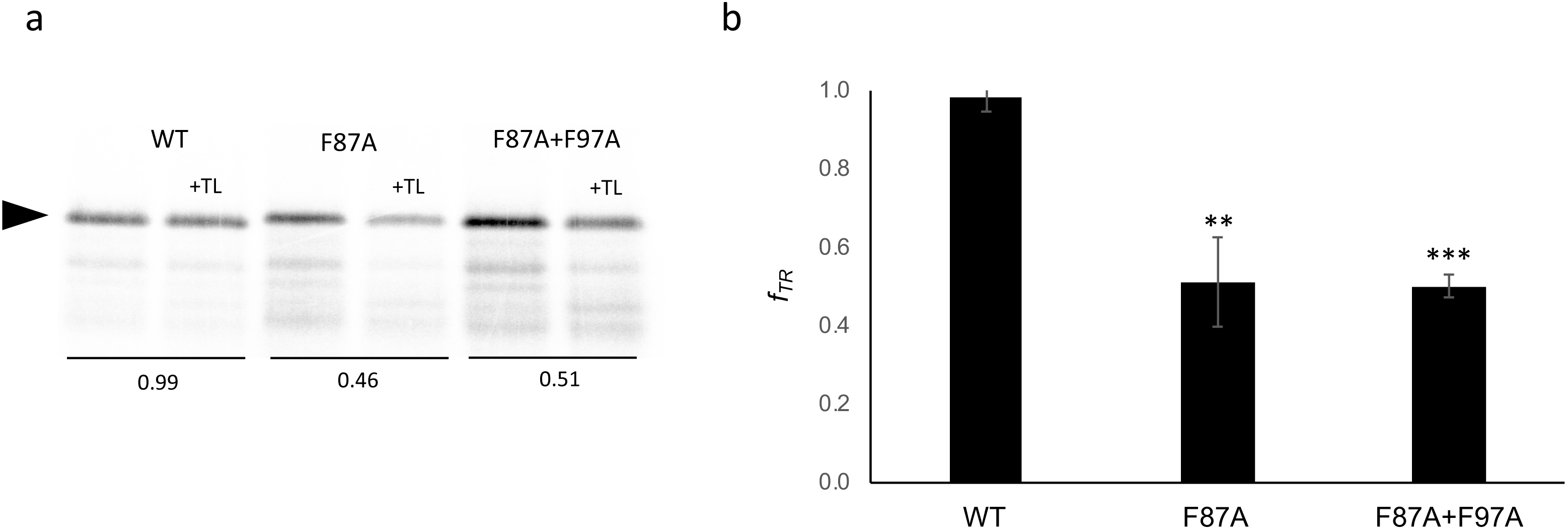
Thermolysin-resistance measurements of PENT (1-217) wildtype and mutants thereof. SDS gel showing the corresponding control and proteolyzed (+TL) products. The black arrow indicates the full-length construct (including an arrest peptide that terminates in a stop codon). The thermolysin-resistant fraction *f_TR_* = *I_TR_* / *I_Buff_,* where *I_TR_* is the intensity of the band after pulse proteolysis and *I_Buff_* is the intensity of the non-treated control band. (b) Quantitation of the data in panel *a*. All experiments were repeated at least 3 times. Averages ± SE are shown, * *p* ≤ 0.05, ** p ≤ 0.01, *** *p* ≤ 0.001.

**Supplementary Table 1.**
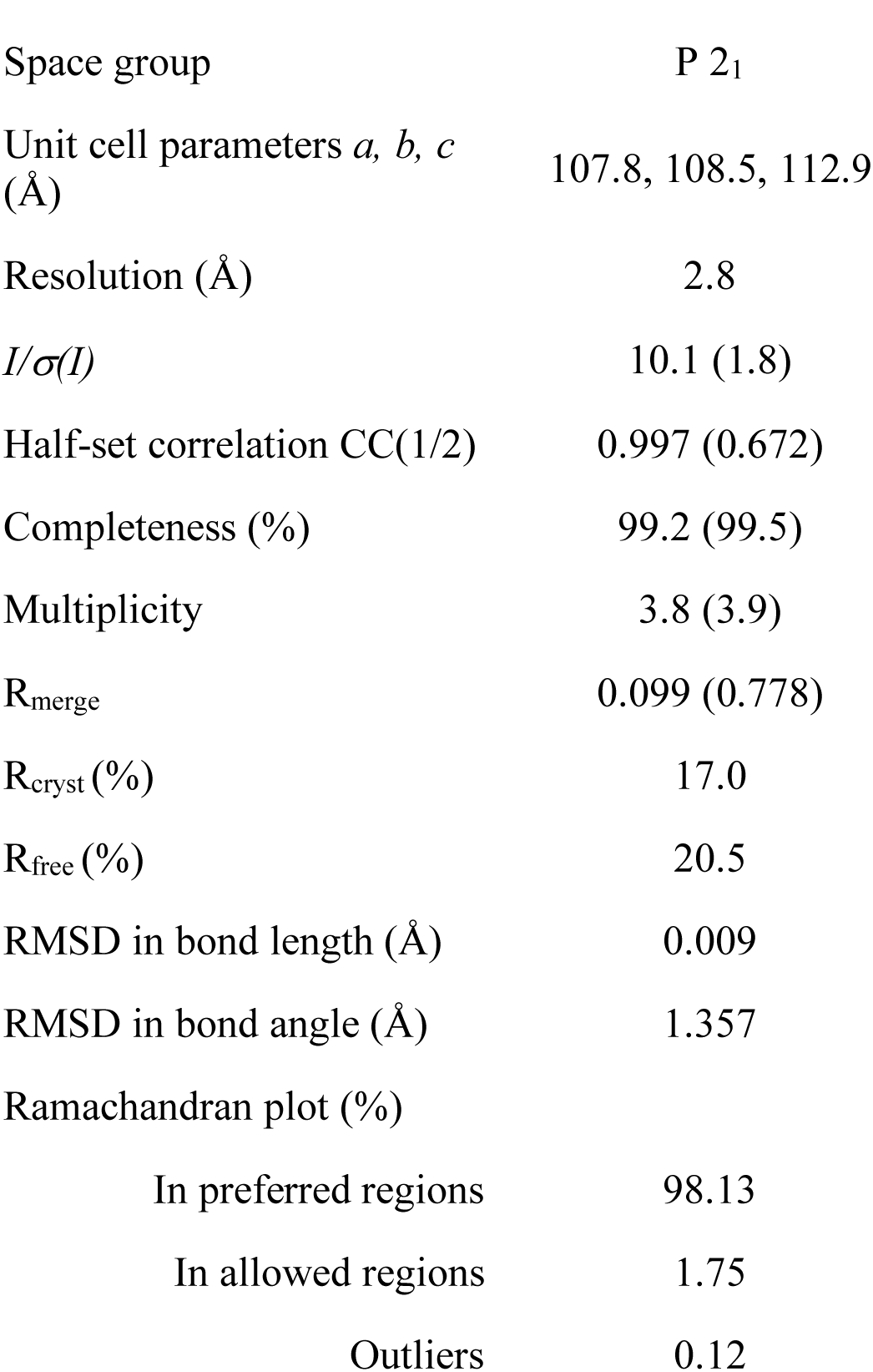
Data processing and refinement statistics for PENT (PDB access code 6FLS).

**Supplementary Table 2.**
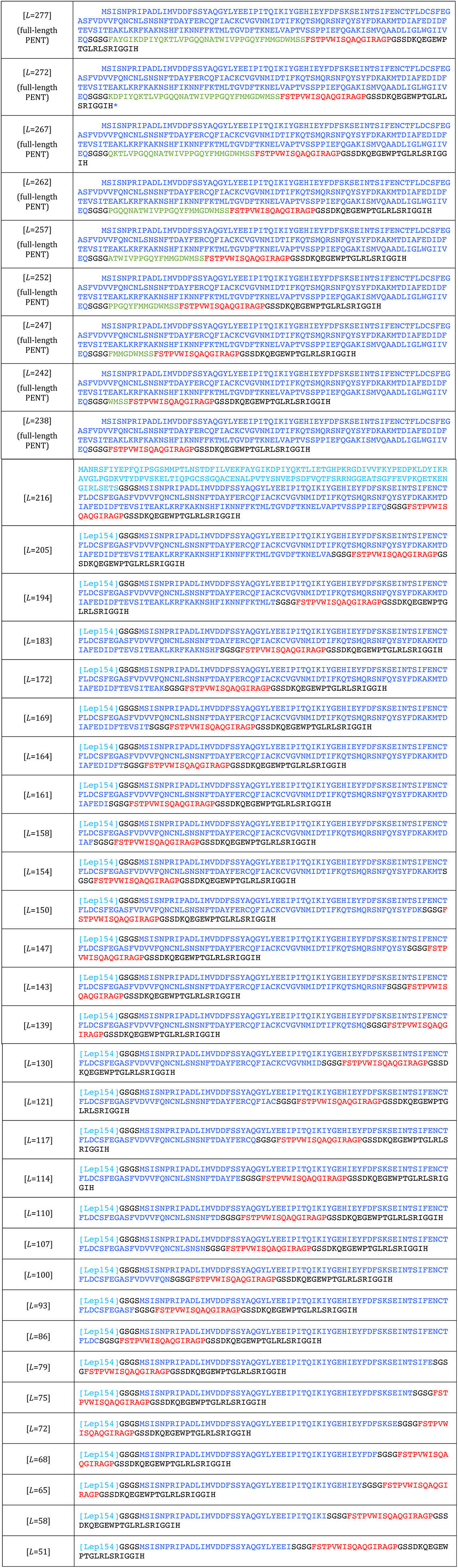
Amino acid sequences of the full-length [Lep154]–PENT–linker–SecM–C-terminal tail construct and of the truncations analyzed in the paper. Note that the N-terminal Lep-part is not present in constructs with *L* ≥ 238 residues (that all contain the full 217-residue PENT domain).

